# Should I stay or should I go? Generalized marginal value theorem with temporal discounting

**DOI:** 10.1101/2024.10.28.620618

**Authors:** Joel Zylberberg

**Affiliations:** Department of Biology, York University, Toronto, Canada; Learning in Machines and Brains Program, Canadian Institute for Advanced Research, Toronto, Canada

**Keywords:** Temporal discounting, Patch foraging, Decision-making, Behavioral economics, Cognitive science, Behavioral ecology, Marginal value theorem, Optimal giving-up times

## Abstract

Consider a person in an environment containing patchy rewards. Under what circumstances should they stay in a given reward patch or leave that patch to seek out a new one? In a landmark 1976 study, Charnov derived the action policy that maximizes the expected rate at which this person gathers rewards. This result is called the Marginal Value Theorem (MVT), and decades of study have shown that humans’ and other animals’ behaviors qualitatively follow MVT, but with notable and systematic deviations. These deviations have been hypothesized to arise from the fact that MVT does not incorporate temporal discounting whereas humans and other animals tend to value current rewards more than future ones. Rigorously testing that hypothesis has been challenging because there is no mathematical theory that determines optimal patch foraging decision policies for agents who use temporal discounting. To fill this knowledge gap, I derived the optimal patch foraging policy for agents who exponentially discount future rewards and studied how that optimal policy depends upon their temporal discount rate, and upon the structure of their environment. Notably there are conditions under which the optimal policy with temporal discounting is to leave *earlier* than is predicted by MVT (under-staying), while under other conditions the optimal policy is to stay longer than is predicted by MVT (over-staying). The theory presented here delineates when each situation arises and may help to interpret the otherwise-puzzling ways in which human and animal behaviors deviate from MVT.

## 1 Introduction

At many of life’s junctures, people face the question of whether to stay in their current situation or to leave and seek out a new one. The ubiquity of this question has motivated many efforts to understand how an idealized optimal agent might address this question, and to understand how and why people and other animals deviate from that optimal behavior. In a landmark 1976 study, Charnov [1] attempted to address this challenge by identifying the decision policy that maximizes the expected rate at which an agent gathers rewards in environments where those rewards are patchy. (For concreteness, one might choose to consider a person foraging for berries where those berries are clustered onto bushes (Fig. 1). In that case, the person must decide when to leave their berry bush to go find a new one.) Importantly, this *patch foraging problem* itself is very general, and the same mathematical framework has been applied to myriad human and animal behaviors. Charnov’s result, called the Marginal Value Theorem (MVT) states that individuals should stay in a patch until their reward rate drops below the expected mean reward rate of their environment, and then leave. This result is deeply appealing for its simplicity, and it has inspired numerous subsequent studies of human and animal behavior [2], including: the gathering of tool-making stones by ancient humans [3, 4]; mushroom gathering [5] and fishing [6] behaviors of modern humans; decisions by hunter-gatherer communities about when to relocate their camps [7]; decisions by non-human primates about when to find new conspecifics with whom to socialize [8]; and patterns of visual fixation points sampled by people during a visual search task [9]. I focus here on studies of human behavior, but note that the same insights apply very broadly to other animals [10, 11] and even to plants [12].

**Fig. 1.**
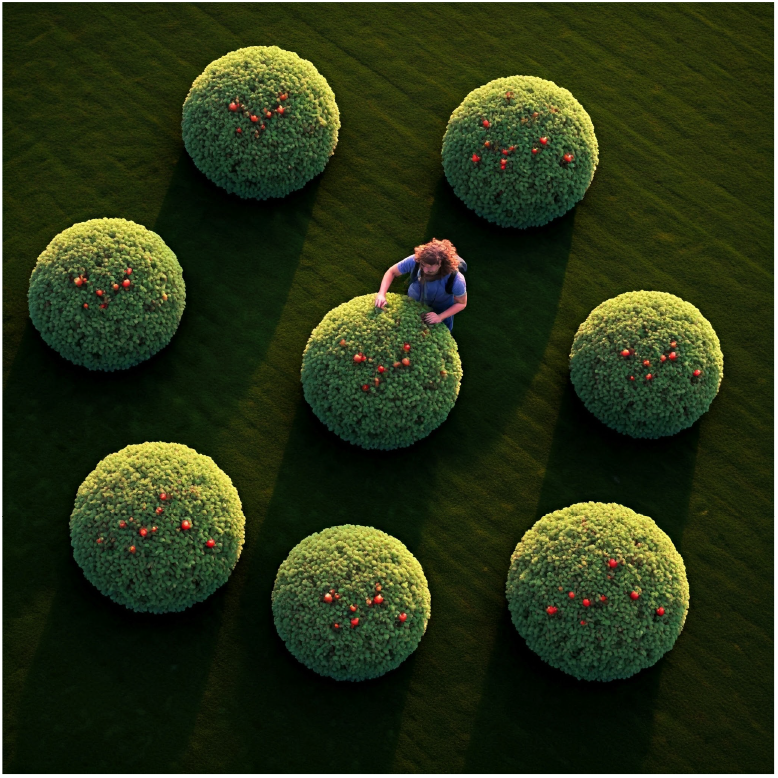
The patch foraging problem. Schematic diagram showing a person foraging within a berry patch. In the distance, there are other berry patches. At each moment, the person must decide whether to stay and harvest berries in their current patch, or to leave and find a new patch in which to forage. (Image generated using the Gemini AI model.)

Across these studies, individuals are typically found to qualitatively follow the predictions of MVT, but with systematic deviations that depend on the experimental condition. Under some conditions, they stay in foraging patches longer than MVT would predict [13, 14], whereas in others, they leave foraging patches earlier than MVT would predict [6, 10]. These deviations have been hypothesized to reflect the fact that individuals have imperfect information about their environment [6], and/or the fact that they value current rewards more than future ones [13]. Identifying the origin of these discrepancies is hampered by the fact that *temporal discounting* was not incorporated into Charnov’s original theory, which means that it is unclear whether and why an observed behavior reflects an optimal temporally discounted action policy. This is an important gap in our knowledge because of how ubiquitously people and other animals are observed to engage in temporal discounting [15, 16], and because of the individual variation in people’s temporal discount rates [17]. The current study aims to fill that gap by deriving a generalized version of MVT that accounts for temporal discounting, and asking when and why the generalized theory predicts over-staying vs under-staying relative to the original undiscounted MVT.

Encouraging this effort, a recent study investigated the behavior of artificial intelligence (AI) agents that had had learned to forage through reinforcement learning (RL) with varying temporal discount rates [18]. That study showed that the AI agents tended to over-stay more when they had larger temporal discount rates; the AI agents were never observed to under-stay relative to the MVT predictions. At the same time, it is difficult to draw firm conclusions about optimal behavior from this AI study for two main reasons: 1) it is possible that the AI agents failed to find the *optimal* action policy within their finite training time and using their finite time horizons for estimating future reward value; and 2) it is unclear how the behavior might differ for conditions that differ from those used in the AI agents’ simulated environments. Relatedly, Wajnberg et al. [19] used dynamic programming to derive a variant of MVT in which rewards are accumulated up to a finite time horizon, and then showed that there is more over-staying (relative to MVT) for agents with shorter time horizons. While the issue of finite time horizons is clearly related to temporal discounting, it does not fully address the question of how temporal discounting affects decision-making when the agent does not have a known, and fixed, time horizon. These limitations point towards the need for new theoretical work to determine the optimal action policy for temporally discounted patch foraging.

To fill that knowledge gap, I generalized Charnov’s original theory to include exponential temporal discounting and derived the optimal patch foraging policy for this scenario. Analyzing that policy, I identified conditions under which agents with temporal discounting will over- or under-stay relative to agents with no discounting (as in the original MVT). Moreover, in the limit of low temporal discount rates, I show that the optimal reward achievable and the optimal decision policy are hyperbolic functions of foraging time and inter-patch travel time. This finding may help to reconcile empirical observations that agents in some studies act consistently with hyperbolic discounting [20, 21], while in others they appear to follow exponential discounting [22, 23]. Finally, I build on the notion that people’s temporal discount rates track expected near-term mortality rates [16] to understand how people’s decisions might be expected to depend on their life circumstances, such as their age, health, and the level of stability within their environment. Collectively, this work helps to understand how and why people behave the way the do. Moreover, it may provide a mathematical and conceptual framework to help individuals make informed decisions at challenging junctures.

## 2 Results

I studied a mathematical model similar to the original one by Charnov [1], but with exponential discounting of future rewards, and with fewer assumptions about how the reward patches change as the person forages within them. The model is as follows.

The environment is assumed to have many different reward patches of varying quality (Fig. 1). Each patch is defined by the reward function *R*_*i*_(*t*), which specifies the time-dependent rate at which the person gathers rewards when they are within that patch. These rates are time dependent because the rewards available in the patch will generally change as the person forages (e.g., the rewards may get depleted as a result of foraging, in which case *R*_*i*_(*t*) will be a decreasing function of time). The person incurs a cost per unit of time spent foraging, which will in general vary between patches. The person is assumed to start in a randomly-chosen reward patch and to continue foraging in that patch for a duration *T*_*i*_, after which they leave the patch to find a new one. They then move on and find a new patch in which to forage. The moving process takes an increment of time and incurs a moving cost: these values are assumed to vary between relocation events. The relevant mathematical symbols are defined in Table 1. Given this scenario, I derived the total exponentially-discounted future reward the person will receive out to an infinite time horizon, as a function of the length of time they spend foraging in each patch, {*T*_*i*_} (Eq. 4 of Section 4.1). I then optimized that total reward with respect to these foraging bout durations, so as to identify the action policy that maximizes the person’s reward (Eq. 5 of Section 4.1).

**Table 1.** Mathematical symbols used in this work.

| Mathematical Symbol | Definition | Comment |
| --- | --- | --- |
| $P_i$ | Probability of patch type $i$ | |
| $t$ | Time | |
| $\lambda$ | Temporal discount rate | |
| $R_i(t)$ | Reward rate at time $t$ in patch type $i$ | |
| $C_{F,i}$ | Foraging cost per unit time in patch type $i$ | |
| $C_M$ | Cost per unit time to move to a new patch | varies between transits |
| $T_M$ | Transit time to move to a new patch | varies between transits |
| $T_i$ | Time spent foraging in patch type $i$ | chosen to maximize $EV(\{T_i\})$ |
| $EV(\{T_i\})$ | Expected total discounted future value | depends on $\{T_i\}$ |
| $V_i(\{T_i\})$ | Discounted future value while in patch type $i$ | $\sum_i P_i V_i(\{T_i\}) = EV(\{T_i\})$ |
| $ER_i(\{T_i\})$ | Expected undiscounted reward rate | |

This derivation is outlined in the Methods Sec. 4.1, and the result represents a generalized form of the marginal value theorem that applies to all temporal discount rates (*λ*), and to any set of patch reward rate functions, foraging costs, travel times, and travel costs. In many cases, the most general result will not be needed because the temporal discount rate *λ* is typically small, and in the case of small *λ*, the mathematical result can be greatly simplified. For that reason, I next analyzed these equations in the limit of small temporal discount rates, *λ*, and obtained the simpler result in Section 4.2, which is as follows:

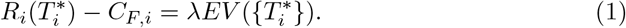

This says that the optimal time at which to leave patch 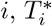, arises when the reward rate (net of foraging costs *C*_*F,i*_) is equal to the expected total future reward 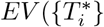, times the temporal discount rate *λ*. This is conceptually similar to Charnov’s original result in that the optimal time to move is when the reward rate matches that which is expected from the overall environment. In the generalized theory which includes temporal discounting, the effective reward rate of the environment is given by the total future reward multiplied by the temporal discount rate *λ*. Importantly, the expected discounted reward 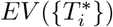 increases as the discount rate *λ* decreases.

In the limit that *λ →* 0, the total expected future reward (out to infinite time horizon), 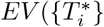, is infinite. Nevertheless, the product 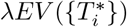 remains finite and is equal to the expected rate of reward acquisition, in exact agreement with Charnov’s MVT result (see Methods Section 4.2). This reinforces the fact that Charnov’s undiscounted marginal value theorem is a special case of the more general theory that I present here. Importantly, in the presence of temporal discounting, the optimal behavior can substantially deviate from that which is expected in the absence of temporal discounting.

Armed with this general theory, I next investigated how the optimal decision policy varies between environments with different characteristics. This analysis enabled me to identify situations in which the optimal agent will either under-stay or over-stay, when compared to an agent who has no temporal discounting. The general theory presented here can be applied to essentially any other situation, and I have publicly shared computer code that can be used to explore different parameter choices and environments (see Methods).

### 2.1 Temporal discounting leads to *over-staying* when the patches are continuously depleted by foraging

While the mathematical theory is very general, and applies to environments with many different types of patches, I analyze here a simplified scenario in which there is only a single type of patch, and that patch type has a reward function that decreases monotonically during the person’s foraging bout (Fig. 2A). This is the scenario originally considered by Charnov [1], and it models an environment where the act of foraging depletes the patch. (E.g., consider a person picking berries from a bush. The number of available berries decreases as they get harvested. Eventually, the person must go find a new bush if they wish to continue picking berries.) Given this scenario, I computed the total discounted expected reward as a function of the foraging bout duration (Eq. 4 of Section 4.1, and Fig. 2B) and determined the bout duration *T* ^*\**^ that maximizes this reward. This optimal bout duration increases with increasing temporal discount rate (Fig. 2C), indicating that optimal agents in this environment who temporally discount will over-stay relative to agents with no temporal discounting.

**Fig. 2.**
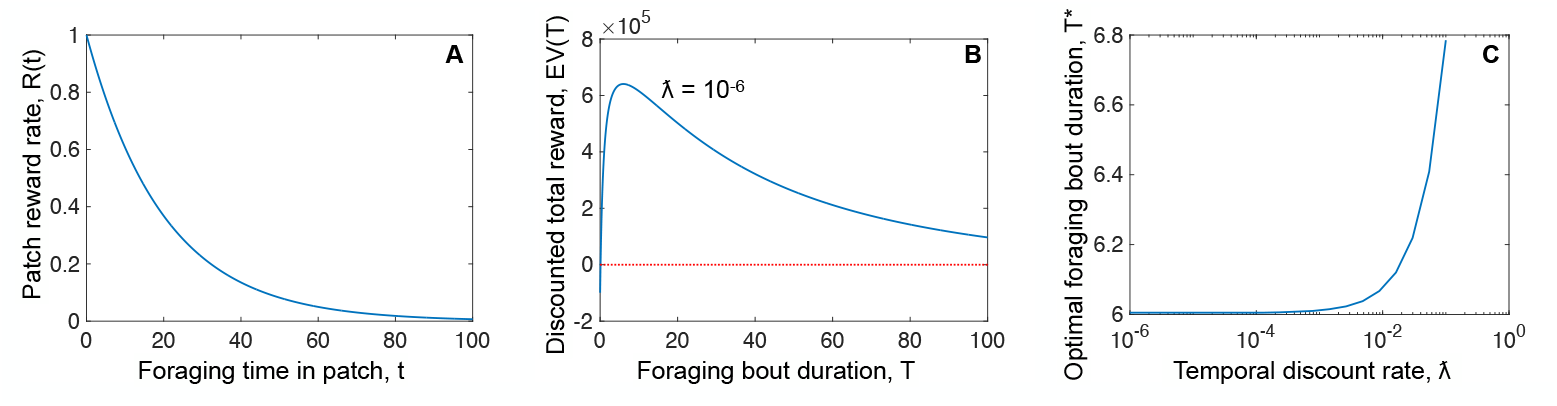
Optimal agents who temporally discount over-stay in an environment with a monotonically decreasing reward rate. (**A**) Rate *R*(*t*) at which an agent in this environment obtains rewards while they are foraging in a patch. (**B**) Total expected temporally discounted reward obtained by the agent (out to an infinite time horizon) as a function of how long they stay in the patch before leaving to find another one. This curve is shown for temporal discount rate *λ* = 10^*−*6^. For larger *λ* values, the curve has a lower amplitude and the peak shifts towards the right. (**C**) Optimal (maximizing *EV* (*T*)) foraging bout duration *T* ^*\**^, as a function of the temporal discounting rate. Optimal agents with larger temporal discount rates stay longer in the patch before leaving to find a new one.

Intuitively, this over-staying can be understood by noting that a person foraging in a patch will continue to get some rewards right away if they stay in that patch, whereas if they leave the patch, their future rewards will be delayed until they get to a new foraging patch. For people who exhibit temporal discounting, those future rewards are worth less than are the currently-available ones. For that reason, they stay in the current patch even as it becomes ever more depleted.

### 2.2 Temporal discounting can lead to *under-staying* when the patches can improve during a foraging bout

Having shown that temporal discounting can lead optimal agents to stay longer in a foraging patch, I next wondered whether and when the opposite effect could arise. In other words, can temporal discounting ever lead optimal agents to under-stay by leaving a foraging patch earlier than would an optimal agent who does not temporally discount? Building on the intuition behind the over-staying observed in Fig. 2, I hypothesized that temporal discounting might cause under-staying in environments where the reward rate can increase over time. In this case, I reasoned that temporal discounting may cause the later rewards to be insufficiently valuable for the optimal agent to stay within a foraging patch for long enough to obtain them, when they could instead move on to a new patch instead.

To test this hypothesis, I repeated the calculations from Fig. 2, but with a different function for the reward rate *R*(*t*). In this new environment, there is an immediate reward that decays over time, followed by a later phase wherein much larger rewards are obtained (Fig. 3A). I again computed the total discounted expected reward as a function of the foraging bout duration (Fig. 3B) and determined the bout duration *T* ^*\**^ that maximizes this reward for different temporal discount rates *λ* (Fig. 3C). In this case, and in stark contrast to the one presented in Fig. 2, we see that agents with sufficiently strong temporal discounting have dramatically shorter optimal foraging bouts as compared to those agents with no discounting. This effect can be understood by analyzing the curves in Figs. 3A,B. For the agent with very little discounting (*λ* = 10^*−*6^), the second peak in the patch’s reward rate (Fig. 3A) corresponds to a large second peak in their expected temporally discounted total reward (Fig. 3B, green curve). For contrast, for the agent with the higher discount rate of *λ* = 10^*−*1^, the second peak in the patch reward rate corresponds to only a tiny second peak in their expected temporally discounted reward (Fig. 3B, black curve). This is because that second reward phase arrives too late to be of much value to the agent with very strong temporal discounting. As a result, optimal agents who strongly temporally discount leave the patch when the initially-available reward starts to dissipate, and long before the second phase of rewards become available. For contrast, optimal agents who less-strongly temporally discount stay long enough to enjoy the second reward phase (Fig. 3C).

**Fig. 3.**
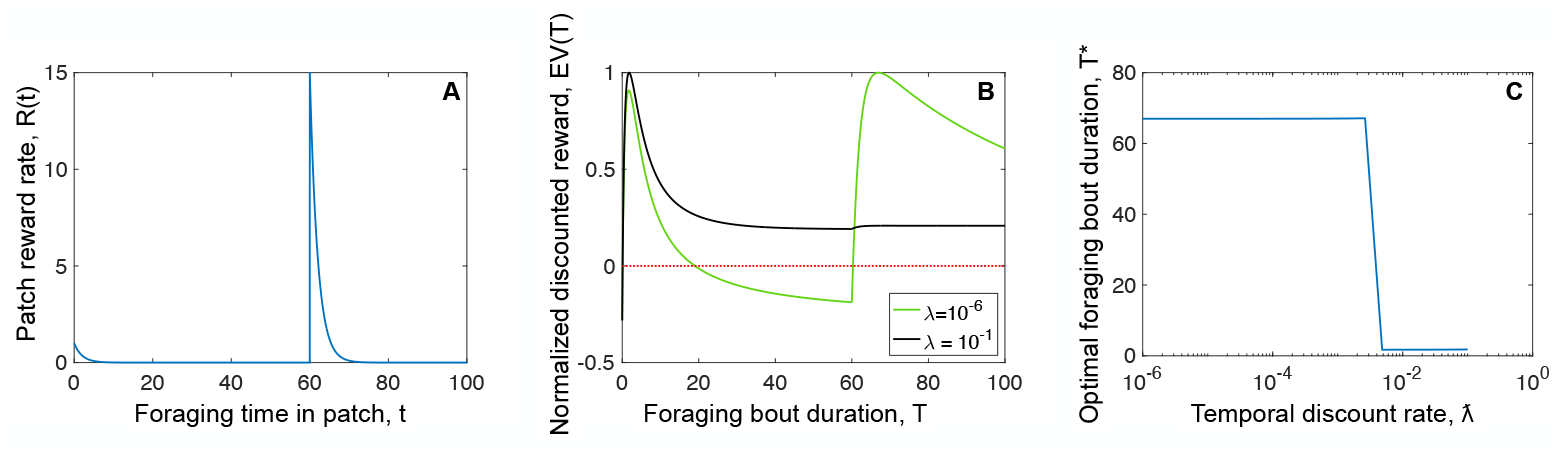
Optimal agents who temporally discount can substantially under-stay in an environment with both early and late rewards. (Similar to Fig. 2 but with a different patch reward rate function *R*(*t*).) (**A**) Rate *R*(*t*) at which an agent in this environment obtains rewards while they are foraging in a patch. A small initial reward decays quickly and is later followed by a much larger reward. (**B**) Total expected temporally discounted reward obtained by the agent (out to an infinite time horizon) as a function of how long they stay in the patch before leaving to find another one. These curves are shown for two different temporal discount rates, and each curve is normalized by its peak amplitude. (**C**) Optimal (maximizing *EV* (*T*)) foraging bout duration *T* ^*\**^, as a function of the temporal discounting rate. Optimal agents with larger temporal discount rates leave the patch much earlier.

## 3 Discussion

People must frequently decide whether to stay in their current situation or to move on and find a new one. E.g., consider the decision to keep fishing in a given spot instead of finding a new one, or to keep picking berries in a given berry patch instead of leaving to find a new one. The classic theoretical framework for understanding how to optimally make these kinds of decisions is called marginal value theorem (MVT) [1] and it does not account for the fact that people and other animals exhibit temporal discounting wherein current rewards are more valuable than are future ones. To overcome this limitation, I derived a generalized version of MVT that incorporates exponential discounting. This generalized theory can be used to understand how optimal decision-making varies between different scenarios, and between people with different temporal discount rates.

Notably, I found that, depending on the environment, temporal discounting can lead optimal agents to either over-stay or under-stay relative to agents who do not temporally discount. This may help to interpret experimental results, where some studies show that individuals over-stay [13, 14], and other studies show that they under-stay [6, 10]. Because optimal agents can exhibit either under-staying or over-staying depending on their discount rate and on the environment, these observed deviations from MVT do not necessarily signify sub-optimal decision-making, nor do they necessarily reflect the individuals having imperfect information with which to make their decisions. This also means that differences between individuals’ decisions in patch foraging type tasks should not necessarily be used to make judgments about the quality of their decision-making: people with different temporal discount rates may make different decisions and all of those decisions may be optimal given the individuals’ discount rates. Indeed, there is substantial individual variation in temporal discount rates [17].

This dependence of optimal decision-making on temporal discount rate is especially interesting given that peoples’ temporal discount rates appear to track their expected near-term mortality rate [16]. In other words, people who are nearer to death exhibit stronger temporal discounting. Coupled with the theoretical work presented here, this means that optimal decisions could be quite different for individuals at different life stages, and in environments with different levels of safety and stability. The theory presented here provides a framework for understanding those differences.

Interestingly, the expected discounted future reward function (Eq. 8) shows a hyperbolic dependence on the transit times and the durations of the foraging bouts. This arises even though the agents being modeled here *exponentially* discount future rewards. This observation – that hyperbolic discounting functions can arise in the small-*λ* limit of exponential discounting – may help to reconcile the observations that people in some studies appear to use exponential discounting [22, 23], while in others, they appear to use hyperbolic discounting [20, 21]. As I observed here, hyperbolic forms of discounting can arise in agents that use exponential discounting, so these two types of temporal discounting may not be as distinct as they first appear.

While the specific examples presented here (i.e., in Fig. 2 and Fig. 3) cannot cover every possible scenario, the mathematical theory itself (Sec. 4.1) is very general. For this reason, I have shared a computer script that can be modified to repeat the analysis in Figs. 2, 3 for different scenarios. I hope and anticipate that others might use this to help interpret their experimental findings on human and animal behavior.

A related theoretical study considered the case of non-consumable rewards within each patch [24]. That work simulated the scenario where the reward functions *R*_*i*_(*t*) are constant in time, and studied how the agents’ propensity to leave their foraging patches depended on the number of other agents present with whom they are competing for resources. The present work does not include the effects of this inter-agent competition. An interesting direction for future work would be to revisit the calculations here, but in the multi-agent setting of Liang and Brinkman [24].

Finally, it is important to note that the mathematics of the patch foraging problem studied here can be applied very generally to any scenario in which a person currently receives rewards (possibly negative ones) and has the option to leave their reward situation and seek out another one. Obvious examples include foraging activities like fishing and gathering berries, but this same framework could be used to understand optimal decisions about jobs (e.g., should I go find a new job or stay in my current one?), monogamous relationships, scientific areas of study, strategic directions in organizations like businesses, moving house, etc. As such, I hope that this normative mathematical framework may be useful beyond interpreting psychology and animal behavior experiments. Rather, I hope and anticipate that it may also be used to help people make otherwise-difficult decisions and to do so in a rigorous and optimal manner.

## 4 Methods

I first present the details of the mathematical calculations, followed by the details for the specific examples shown in the Results section. The mathematical symbols used in this work are defined in Table 1.

### 4.1 Mathematical Derivation of the Temporally Discounted Marginal Value Theorem

Let us first calculate the expected discounted future reward value, *V*_*i*_({*T*_*i*_}), for an agent that has just arrived in a patch of type *i* at time *t* = 0. That expected reward depends on how long the agent stays in this patch, and on how long they will stay in all future patches they encounter. Thus, the expected discounted future reward depends on the foraging durations in *all* patch types.

That calculation proceeds as follows. First, I calculate the accumulated discounted reward over three different epochs: 1) the epoch during which the agent forages in the current patch; 2) the travel epoch – of random duration – after they leave the current patch in search of a new patch; and 3) the entire future – out to infinite time horizon – after they arrive at the next patch. I then sum these accumulated discounted rewards to obtain

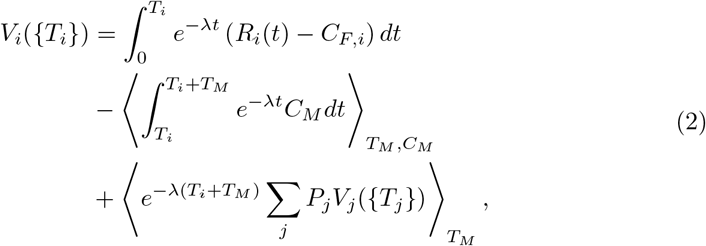

where the angular brackets denote an expectation value over the distribution of travel costs *C*_*M*_ and/or travel times *T*_*M*_, as noted in the subscripts.

I next calculate the expectation value of these rewards over various the types patches in which the agent could currently be located:

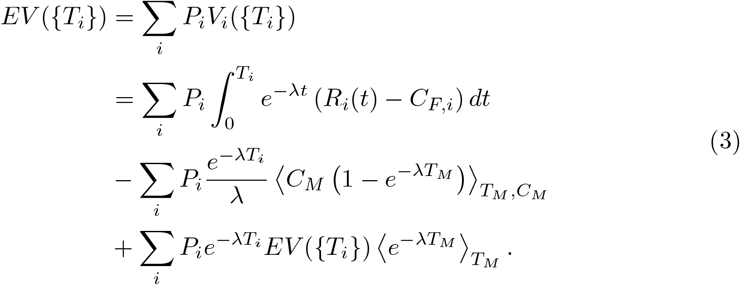

Note that this equation has a very convenient recursive structure which will enable us to compute the total future discounted reward, up to an infinite time horizon. This is similar to the reasoning used in our previous studies of prey animal decision-making [25] and image statistics [26], and also resembles the logic that underlies Bellman’s work in dynamic programming [27]. Subtracting 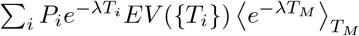 from both sides of Eq. 3, and dividing both sides by 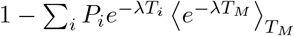, we obtain

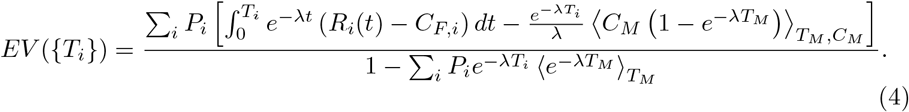

Our next task is to optimize Eq. 4 with respect to the durations of the foraging bouts in each patch type, {*T*_*i*_}. This involves some straightforward but tedious calculus wherein I calculate 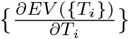, and determine the optimality condition for which all of these partial derivatives vanish. The result is

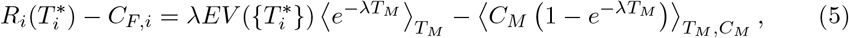

where 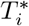 denotes the optimal foraging bout duration in patch *i*, and all of the other variables in Eq. 5 are the independent variables of the problem. The left hand side of the second line of Eq. 5 is the rate at which rewards are obtained from patch *i* (net of foraging costs) at the end point of the optimal-duration foraging bout within that patch. The right hand side of the equation depends on the total expected future reward for this optimal policy, 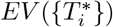, in addition to the moving cost, the transit time, and the discount rate. Note that 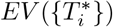 depends on the temporal discount rate, *λ*, although the notation is simplified so as to not list all of the variables upon which 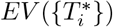 depends.

Together, Eqs. 4 and 5 represent a generalized form of the marginal value theorem, that makes no assumptions about the magnitude of the temporal discounting, and applies to stochastic patch reward rates, foraging costs, travel times, and travel costs. While this result is very general, it is also somewhat complicated. Below, I will analyse this result for the case of small temporal discount rates. This will enable us to greatly simplify Eqs. 4 and 5, enabling us to develop deeper intuition for the problem. It will also enable us to reproduce Charnov’s original MVT result, which is obtained from our generalized result in the limit as *λ* → 0.

Note that, if we simply evaluate Eq. 4 for the *λ* = 0 case of no temporal discounting, we find that the denominator vanishes and the expected total future reward is infinite. (This makes sense: summing future rewards over an infinite time horizon without discounting yields infinite total reward.) Despite this complication, the product *λEV* ({*T* ^*\**^}) remains finite as *λ* → 0, which will enable us to analyze the optimal decision criterion in Eq. 5 in this small-*λ* case, and to recover Charnov’s original result for the special case of *λ* → 0. I will return to this issue later, after first deriving the small-*λ* approximations to Eqs. 4 and 5.

### 4.2 Simplifying the Result in the Case of Small Temporal Discount Rates

So long as 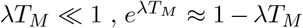. This approximation holds for small *λ*. In that case, Eq. 5 simplifies to

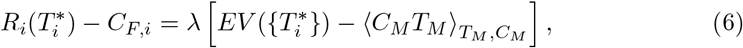

where I have omitted terms of second order and higher in *λ*. As I previously noted, 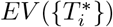 is very large in the case of small *λ*. This means that, for sufficiently small *λ* and finite 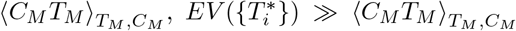 and thus 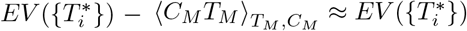. Consequently, Eq. 6 simplifies to

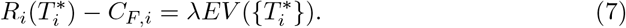

It is important to recall that 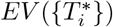 is the expected *total* discounted future reward, and not the *rate* at which rewards are acquired.

To proceed further in simplifying our analysis, I evaluate *λEV* ({*T*_*i*_}) in the small *λ* limit, using Eq. 4 and the approximation 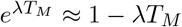 :

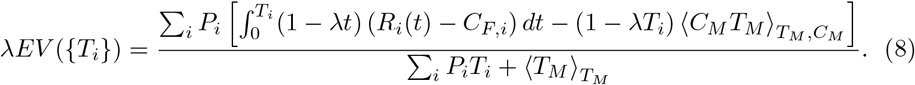

In the limit that *λ* → 0, this becomes

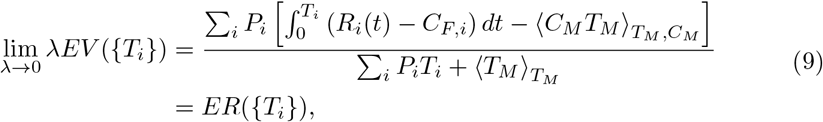

where *ER*({*T*_*i*_}) is the undiscounted expected reward rate of the environment. (The fraction on the right hand side of Eq. 9 is the ratio between the expected reward accumulated during a foraging bout in one patch including the travel period to find the next patch, and the expected amount of time spent on that foraging bout and transit period. It is thus the expected reward rate.) Thus, in this limit, the criterion that defines the optimal behavior (Eq. 7) becomes

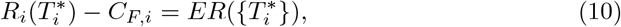

which is the same result as in Charnov’s 1976 study [1].

### 4.3 Details for the specific examples presented herein

For the examples in Figs. 2 and 3, I wrote a short computer script that defines all of the quantities in Table 1 and then calculates both the total discounted future reward, and the optimal foraging bout duration. That code is freely available at the following link and I hope that others will use it to investigate scenarios beyond those that were considered here: http://www.jzlab.org/Plot MVT Curves 1patch GeneralRofT.m.

For the examples in both Figs. 2 and 3, *T*_*M*_ = 1, and *C*_*M*_ = *C*_*F*_ = 0.1. The two examples differ in terms of their reward rate function *R*(*t*). For Fig. 2, *R*(*t*) = *e*^*−*0.05*t*^ whereas for Fig. 3, *R*(*t*) = *e*^*−*0.5*t*^ + 15Θ(*t* − 60)*e*^*−*0.5(*t−*60)^, where Θ denotes the Heaviside step function.

## Acknowledgements

Thanks to Craig Chapman and Nathan Wispinski for introducing me to this interesting problem and to Braden Brinkman for helpful feedback. This work was supported by a Canada Research Chair Grant (CRC-2022-00277) to author JZ.

